# NEMO: Cancer subtyping by integration of partial multi-omic data

**DOI:** 10.1101/415224

**Authors:** Nimrod Rappoport, Ron Shamir

## Abstract

**Motivation:** Cancer subtypes were usually defined based on molecular characterization of single omic data. Increasingly, measurements of multiple omic profiles for the same cohort are available. Defining cancer subtypes using multi-omic data may improve our understanding of cancer, and suggest more precise treatment for patients.

**Results:** We present NEMO (NEighborhood based Multi-Omics clustering), a novel algorithm for multiomics clustering. Importantly, NEMO can be applied to partial datasets in which some patients have data for only a subset of the omics, without performing data imputation. In extensive testing on ten cancer datasets spanning 3168 patients, NEMO outperformed nine state-of-the-art multi-omics clustering algorithms on full data and on imputed partial data. On some of the partial data tests, PVC, a multiview algorithm, performed better, but it is limited to two omics and to positive partial data. Finally, we demonstrate the advantage of NEMO in detailed analysis of partial data of AML patients. NEMO is fast and much simpler than existing multi-omics clustering algorithms, and avoids iterative optimization.

**Availability:** Code for NEMO and for reproducing all NEMO results in this paper is in github.

**Supplementary information:** Supplementary data are available online.

## 1 Introduction

Recent technological advances have facilitated the production of multiple genome-wide high throughput biological data types, collectively termed ”omics”. These include genomics, transcriptomics, proteomics and many more. Analysis of omics datasets was proven invaluable for basic biological research and for medicine. Until recently, research in computational biology has focused on analyzing a single omic type. While such inquiry provides insights on its own, methods for integrative analysis of multiple omic types may reveal more holistic, systems-level insights.

Omic profiles of large cohorts collected in recent years can help to better characterize human disease, facilitating more personalized treatment of patients. In oncology, analysis of large datasets has led to the discovery of novel cancer subtypes. The classification of tumors into these subtypes is now used in treatment decisions (Parker *et al*., 2009; Prasad *et al*., 2016). However, these subtypes are usually defined based on a single omic (e.g. gene expression), rather than through an integrative analysis of multiple data types. The large international projects like TCGA (McLendon *et al*., 2008) and ICGC (Zhang *et al*., 2011) now provide multi-omic cohort data, but better methods for their integrated analysis are needed. Novel, improved methods that employ multiple data types for cancer subtyping can allow us to better understand cancer biology, and to suggest more effective and precise therapy (Kumar-Sinha and Chinnaiyan, 2018; Senft *et al*., 2017).

### 1.1 Multi-Omics Clustering Approaches

There are several approaches to multi-omics clustering (see the reviews of Wang and Gu (2016), Huang *et al*. (2017), Rappoport and Shamir (2018)). The simplest approach, termed *early integration*, concatenates all omic matrices and applies single-omic clustering on the resulting matrix. LRAcluster (Wu *et al*., 2015) is an example of such a method, which probabilistically models the distribution of numeric, count and discrete features. Early integration increases the dimensionality of the data, and ignores the different distributions of values in different omics.

*Late integration* methods cluster each omic separately, and then integrate the clustering results, for example using consensus clustering (Monti *et al*., 2003). PINS (Nguyen *et al*., 2017) is a late integration method that defines connectivity matrices as describing the co-clustering of different samples within an omic, and integrates these matrices. Late integration ignores interactions that are weak but consistent across omics.

*Middle integration* approaches build a single model that accounts for all omics. These models include joint dimension reduction of omic matrices and similarity (kernel) based analyses. Dimension reduction approaches include jNMF, MultiNMF (Zhang *et al*., 2012; Liu *et al*., 2013), iCluster (Shen *et al*., 2009), and its extensions iClusterPlus and iClusterBayes (Mo *et al*., 2013, 2018). CCA is a classic dimension reduction algorithm (Hotelling, 1936), which linearly projects two omics to a lower dimension such that the correlation between the projections is maximal. MCCA (Witten and Tibshirani, 2009) generalizes CCA to more than two omics. Because of the high number of features and the complexity of dimension reduction algorithms, feature selection is required. Similarity based methods handle these shortcomings by working with inter-patient-similarities. These methods have improved runtime, and are less reliant on feature selection. Examples are SNF (Wang *et al*., 2014) and rMKL-LPP (Speicher and Pfeifer, 2015). SNF builds a similarity network of patients per omic, and iteratively updates these networks to increase their similarity until they converge to a single network, which is then partitioned using spectral clustering. rMKL-LPP uses dimension reduction, such that similarities between neighboring samples is maintained in lower dimension. For that purpose, it employs multiple kernel learning, using several different kernels per omic, and providing flexibility in the choice of the kernels. All the middle integration methods above use iterative optimization algorithms, and in some cases guarantee only convergence to local optimum.

To the best of our knowledge, to date, all middle integration methods for multi-omics clustering developed within the bioinformatics community assume full datasets, i.e., data from all omics were measured for each patient. However, in real experimental settings, often for some patients only a subset of the omics were measured. We call these *partial datasets* in the rest of the paper. This phenomenon is already prevalent in existing multi-omic datasets, such as TCGA (McLendon *et al*., 2008), and will increase as cohorts grow. Being able to analyze partial data is of paramount importance, due to the high cost of experiments, and the unequal cost for acquiring data for different omics. Naive solutions like using only those patients with all omics measured or imputation have obvious disadvantages.

A close problem to multi-omics clustering was researched in the machine learning community. In the area of ”multi-view learning” (reviewed in Zhao *et al*. 2017), methods for multi-view clustering actually solve the multiomic clustering problem. PVC (Li *et al*., 2014) is such a method for clustering in the presence of partial data, which is based on joint nonnegative matrix factorization, such that the objective function only considers observed values. This method has not been previously applied on multi-omic data.

### 1.2 Our Contribution

We present NEMO (NEighborhood based Multi-Omics clustering), a simple algorithm for multi-omics clustering. NEMO does not require iterative optimization and is faster than prior art. NEMO is inspired and bulids on prior similarity-based multi-omics clustering methods such as SNF and rMKL-LPP. NEMO’s implementation, as well as code to reproduce the results in this paper, are available in github: https://github.com/Shamir-Lab/NEMO.

We evaluated the performance of NEMO by comparing it to a wide range of multi-omics clustering methods on several cancer data types. On full datasets, despite its simplicity, NEMO showed an improvement over leading multi-omics clustering algorithms. In order to evaluate performance on partial multi-omic data, we compared NEMO to PVC and to data imputation followed by clustering using several methods. In most tests on synthetic data and on real cancer data, NEMO had clear advantage. Finally, we analyzed NEMO’s clustering solution for Acute Myeloid Leukemia, and showed the merit of using multiple omics with partial data.

## 2 Methods

NEMO works in three phases. First, an inter-patient similarity matrix is built for each omic. Next, the matrices of different omics are integrated into one matrix. Finally, that network is clustered.

### 2.1 NEMO - full omics datasets

The input to NEMO is a set of data matrices of *n* subjects (samples or pateints). Given *L* omics, let *X*_*l*_ denote the data matrix for omic l. *X*_*l*_ has dimensions *p*_*l*_ x *n*, where *p*_*l*_ is the number of features for omic *l*. *P* = Σ_*l*_*p*_*l*_ is the total number of features.

Denote by *x*_*li*_ the profile of sample *i* in omic *l* (column *i* in *X*_*l*_). Let *η*_*li*_ denote its *k* nearest neighbors within omic *l*, where Euclidean distance is used to measure profile closeness. For omic *l*, an *n* x *n* similarity matrix *S*_*l*_ is defined as follows:

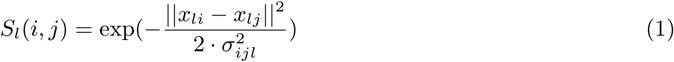

where 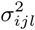 is defined by:

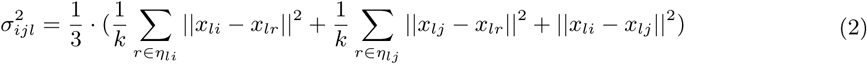

This similarity measure is based on the radial basis function kernel (Buhmann, 2003). 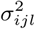 is a normalizing factor, which controls for the density of samples by averaging the distance of the *i*’th and *j*’th samples to their nearest neighbors and the distance between these two samples (Wang *et al*., 2014; Bo Wang *et al*., 2012; Yang *et al*., 2008).

Next, we define the relative similarity matrix, *RS*_*l*_, for each omic:

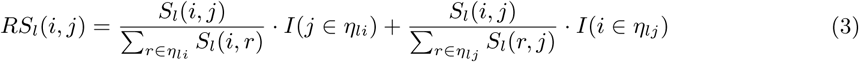

where *I* is the indicator function. *RS*_*l*_(*i, j*) measures the similarity between *i* and *j* relative to *i*’s *k* nearest neighbors and to *j*’s *k* nearest neighbors. Since different omics have different data distributions, the relative similarity is more comparable between omics than the original similarity matrix *S*.

In the next step, NEMO calculates the *n* x *n* average relative similarity matrix *ARS* as:

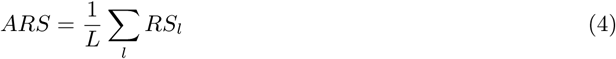

*RS*_*l*_ can be viewed as defining a transition probability between samples, such that the probability to move between samples is proportional to their similarity. Such transition distributions are widely used to describe random walks on graphs (Lo Asz, 1993). *ARS* is therefore a mixture of these distributions (Zhou and Burges, 2007).

Given *ARS*, the clusters are calculated by performing spectral clustering on *ARS* (von Luxburg, 2007). We use the spectral clustering variant that is based on the eigenvectors of the normalized Laplacian, developed by Ng *et al*. (2001).

To determine the number of clusters, we use a modified eigengap method (von Luxburg, 2007). The number of clusters is set to *argmax*_*i*_(*λ*_*i*+1_ - *λ_i_*) · *i*, where *λ* are *ARS* eigenvalues. Intuitively, this objective maintains the idea of the eigengap while encouraging the solution to have a higher number of clusters. This is desired since we observed that often some increase in the number of clusters compared to that prescribed by the eigengap method improved the prognostic value for cancer data. The number of clusters determined by this method is at least as high as the number determined using the eigengap method.

As suggested by Wang *et al*. (2014), we set the number of neighbors in each omic to be the 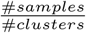 in case the number of clusters is known. When the number of clusters is not known, we use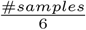. We show NEMO’s robustness to that parameter.

### NEMO - partial datasets

NEMO can handle samples that were measured on only a subset of omics. Specifically, we require that each pair of samples has at least one omic on which they were both measured. Note that this holds in particular if there is an omic for which all samples have measurements, which is often the case for gene expression data. Under these conditions, *RS*_*l*_ is computed as in the full-data scenario, but ARS is now only averaged on the observed values. Denote by *JM* (*i, j*) the omic types available for both samples. Then:

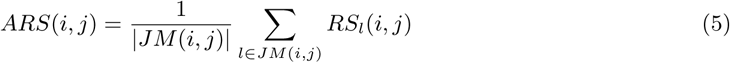

Note that we require that all samples that have measurements for some omic, have measurements for the same set of features in that omic, such that even in the partial data settings each *X*_*l*_ is a full matrix, albeit with fewer rows. For example, the expression of the same set of genes is measured for all patients with RNA-seq data. When patients have different sets of measured features in the same omic, either intersection of the features or imputation of missing values is required.

On partial datasets, each omic *l* may have a different number of samples #*samples*(*l*). The number *k* of nearest neighbors is chosen per omic. Generalizing the full data setting, for omic *l* we set 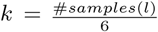

### Time complexity

Computing the distance between a pair of patients in omic *l* takes *O*(*p*_*l*_), so calculating the distance between all patients in all omics takes *O*(*n*^2^ · *P*). The *k* nearest neighbors of each patient and its average distance to them in a specific omic can be computed in time *O*(*n*) per patient (Blum *et al*., 1973), for a total of *O*(*n*^2^ · *L*). Given the distances, the nearest neighbors, and the average distance to them, each 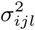 can be computed in *O*(*k*) time. Each entry in *RS*_*l*_ is also calculated in *O*(*k*). *ARS* calculation therefore requires *O*(*n*^2^ · *P*), and spectral clustering takes *O*(*n*^3^), so the total time is *O*(*n*^2^ · *P* + *n*^3^).

Other similarity-based methods such as SNF and rMKL-LPP need the same *O*(*n*^2^*·P*) time to calculate the distances. However, the iterative procedure in both SNF and rMKL-LPP requires *O*(*n*^3^) per iteration.

### Clustering assessment

For the simulated datasets, to gauge the agreement between a clustering solution and the correct cluster structure, we used the adjusted Rand index (ARI) (Hubert and Arabie, 1985).

To assess clustering solutions for real cancer samples, we used survival data and clinical parameters reported in TCGA. We used the logrank test for survival (Hosmer *et al*., 2008) and enrichment tests for clinical parameters. We used the *χ*^2^ test for independence to calculate enrichment of discrete clinical parameters, and Kruskal-Wallis test for numerical parameters. It was previously observed that the *χ*^2^ approximation for the statistic of these tests produces biased p-values that overestimate the significance (Vandin *et al*., 2015; Rappoport and Shamir, 2018). In order to better approximate the real p-values, we performed permutation tests on the clustering solution, and reported the fraction of permutations for which the test statistic was greater or equal to that of the original clustering solution as the empirical p-value. Full details on the permutation testing appear in Rappoport and Shamir (2018).

## Results

We applied NEMO in several settings. First, we compared it to nine multi-omics clustering algorithms on ten *full* cancer datasets. We next compared NEMO to several methods on simulated partial data, and on real cancer datasets with parts of the data artificially removed. Finally, we used NEMO on a real partial cancer dataset.

### Full datasets

We applied NEMO to ten TCGA datasets spanning 3168 patients. The datasets are for the following cancer types: Acute Myeloid Leukemia (AML), Breast Invasive Carcinoma (BIC), Colon Adenocarcinoma (COAD), Glioblastoma Multiforme (GBM), Kidney Renal Clear Cell Carcinoma (KIRC), Liver Hepatocellular Carcinoma (LIHC), Lung Squamous Cell Carcinoma (LUSC), Skim Cutaneous Melanoma (SKCM), Ovarian serous cystadenocarcinoma (OV) and Sarcoma (SARC). For each dataset, we analyzed three omics: gene expression, methylation and miRNA expression. When some of the patients lacked measurements for some of the omics, we included only those patients that had data from all omics.

We have previously used these datasets to benchmark multi-omics clustering methods (Rappoport and Shamir, 2018). Datasets sizes varied between 170 and 621 samples. See Rappoport and Shamir (2018) for full details and preprocessing. Clustering results for NEMO on all datasets are in Supplementary File 2.

We compared NEMO on each dataset to nine different multi-omics clustering methods. As early integration methods we used LRAcluster, and k-means and spectral clustering on the concatenation of all omic matrices. For late integration we used PINS. We used MCCA, MultiNMF and iClusterBayes as joint dimension reduction methods. Finally, SNF and rMKL-LPP represented similarity-based integration. We set *k*, the number of neighbors in NEMO to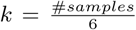. For all methods, we chose the number of clusters in the range 2-15 using the methods recommended by the authors. The results of the nine methods were taken from our benchmark study (Rappoport and Shamir, 2018), where full details on the execution of all methods are available. (For MCCA, LRAcluster and k-means the results are slightly different, since here we increased the number of k-means repeats they perform in order to increase their stability.)

To assess the clustering solutions we compared the survival curves of different clusters, and performed enrichment analysis on clinical labels (see Methods section). To avoid biases, we chose the same set of clinical parameters for all cancers: age at initial dianosis, gender, and four discrete clinical pathological parameters. These parameters quantify the progression of the tumor (pathologic T), cancer in lymph nodes (pathologic N), metastases (pathologic M) and total progression (pathologic stage). In each cancer type we tested the enrichment of each parameter that was available for it.

Table 1, Figure 1 and Table 2 summarize the performance of the ten algorithms on the ten datasets. NEMO found a clustering with significant difference in survival for six out of ten cancer types, while all other methods found at most five. None of the methods found a clustering with significantly different survival for the COAD, LUSC and OV datasets. The p-value for KIRC, the only other dataset for which NEMO did not reach significance, was 0.063. NEMO had an average logrank p-value of 1.64, second after MCCA (1.75). NEMO found at least one enriched clinical parameter in eight of the ten datasets, the highest number found and tied with spectral clustering, rMKL-LPP and MCCA. The average number of enriched clinical parameters for NEMO was 1.5, second only to rMKL-LPP with 1.6.

**Table 1:** Aggregate statistics of the tested multi-omics clustering methods across ten cancer datasets. First row: number of datasets with significantly different survival. Second row: number of datasets with at least one enriched clinical label. Third row: mean number of clusters. Fourth row: mean runtime. Best performers in each category are marked in bold.

|  | K-means | Spectral | LRAcluster | PINS | SNF | rMKL-LPP | MCCA | MultiNMF | iClusterBayes | NEMO |
| --- | --- | --- | --- | --- | --- | --- | --- | --- | --- | --- |
| Significantly different survival | 2 | 3 | 5 | 4 | 4 | 4 | 5 | 4 | 2 | **6** |
| Significant clinical enrichment | 6 | **8** | 7 | 6 | 6 | **8** | **8** | 6 | 5 | **8** |
| Number of clusters | 2.6 | 3.3 | 9.4 | 4.6 | 2.8 | 6.7 | 10.6 | 2.2 | 2.2 | 4.5 |
| Runtime (sec) | 394 | **4** | 3388 | 147 | 13 | 233 | 20 | 28 | 4298 | 10 |

**Table 2:**
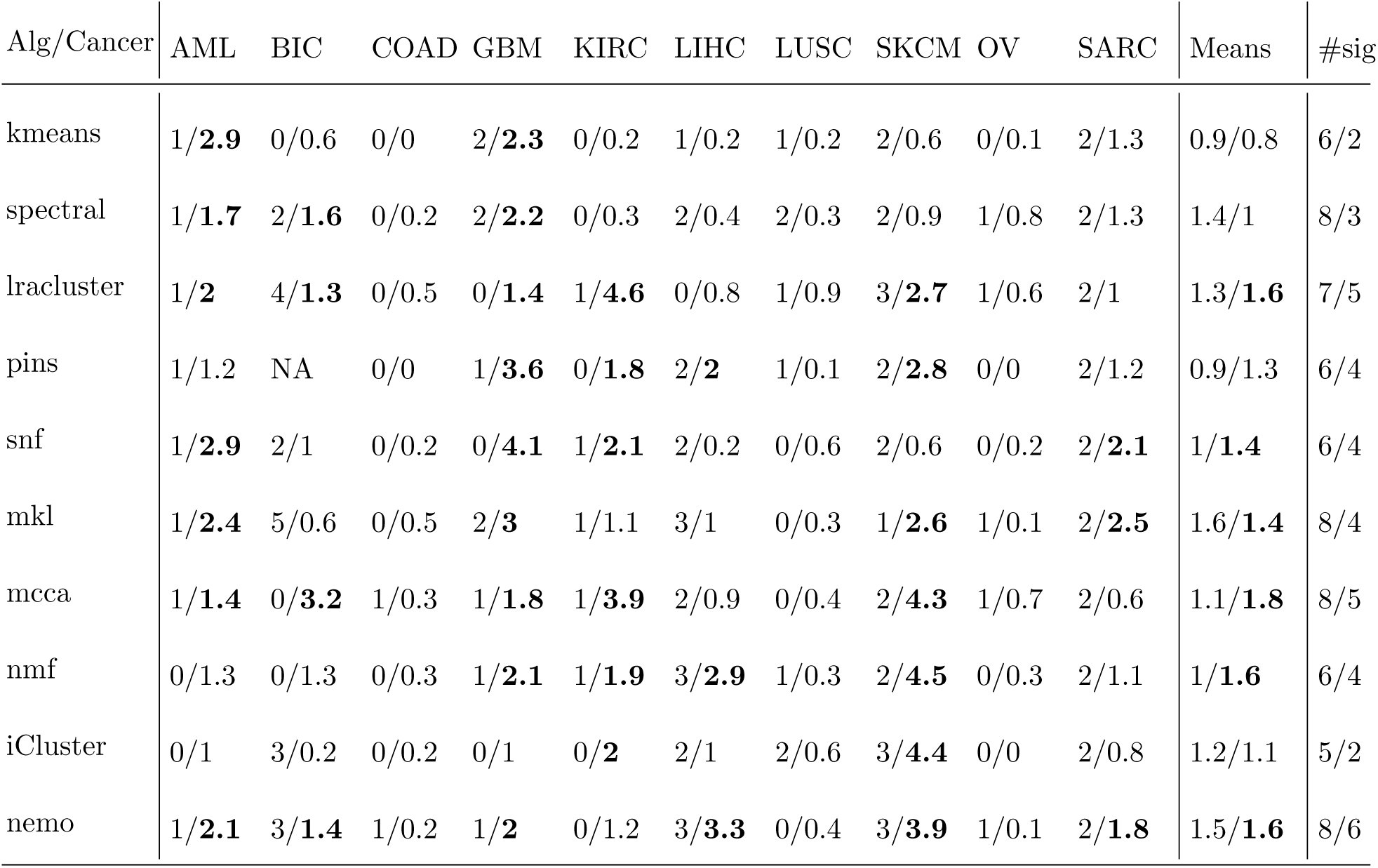
Results of applying the ten algorithms on cancer datasets. The first number in each cell is the number of significant clinical parameters detected, and the second number is the -log10 p-value for survival, with bold numbers indicating significant results. Means are algorithm averages. #sig is the number of datasets with significant clinical/survival results.

| Alg/Cancer | AML | BIC | COAD | GBM | KIRC | LIHC | LUSC | SKCM | OV | SARC | Means | #sig |
| --- | --- | --- | --- | --- | --- | --- | --- | --- | --- | --- | --- | --- |
| kmeans | 1/**2.9** | 0/0.6 | 0/0 | 2/**2.3** | 0/0.2 | 1/0.2 | 1/0.2 | 2/0.6 | 0/0.1 | 2/1.3 | 0.9/0.8 | 6/2 |
| spectral | 1/**1.7** | 2/**1.6** | 0/0.2 | 2/**2.2** | 0/0.3 | 2/0.4 | 2/0.3 | 2/0.9 | 1/0.8 | 2/1.3 | 1.4/1 | 8/3 |
| lrcluster | 1/**2** | 4/**1.3** | 0/0.5 | 0/**1.4** | 1/**4.6** | 0/0.8 | 1/0.9 | 3/**2.7** | 1/0.6 | 2/1 | 1.3/**1.6** | 7/5 |
| pins | 1/1.2 | NA | 0/0 | 1/**3.6** | 0/**1.8** | 2/**2** | 1/0.1 | 2/**2.8** | 0/0 | 2/1.2 | 0.9/1.3 | 6/4 |
| snf | 1/**2.9** | 2/1 | 0/0.2 | 0/**4.1** | 1/**2.1** | 2/0.2 | 0/0.6 | 2/0.6 | 0/0.2 | 2/**2.1** | 1/**1.4** | 6/4 |
| mkl | 1/**2.4** | 5/0.6 | 0/0.5 | 2/**3** | 1/1.1 | 3/1 | 0/0.3 | 1/**2.6** | 1/0.1 | 2/**2.5** | 1.6/**1.4** | 8/4 |
| mcca | 1/**1.4** | 0/**3.2** | 1/0.3 | 1/**1.8** | 1/**3.9** | 2/0.9 | 0/0.4 | 2/**4.3** | 1/0.7 | 2/0.6 | 1.1/**1.8** | 8/5 |
| nmf | 0/1.3 | 0/1.3 | 0/0.3 | 1/**2.1** | 1/**1.9** | 3/**2.9** | 1/0.3 | 2/**4.5** | 0/0.3 | 2/1.1 | 1/**1.6** | 6/4 |
| iCluster | 0/1 | 3/0.2 | 0/0.2 | 0/1 | 0/**2** | 2/1 | 2/0.6 | 3/**4.4** | 0/0 | 2/0.8 | 1.2/1.1 | 5/2 |
| nemo | 1/**2.1** | 3/**1.4** | 1/0.2 | 1/**2** | 0/1.2 | 3/**3.3** | 0/0.4 | 3/**3.9** | 1/0.1 | 2/**1.8** | 1.5/**1.6** | 8/6 |

**Figure 1:**
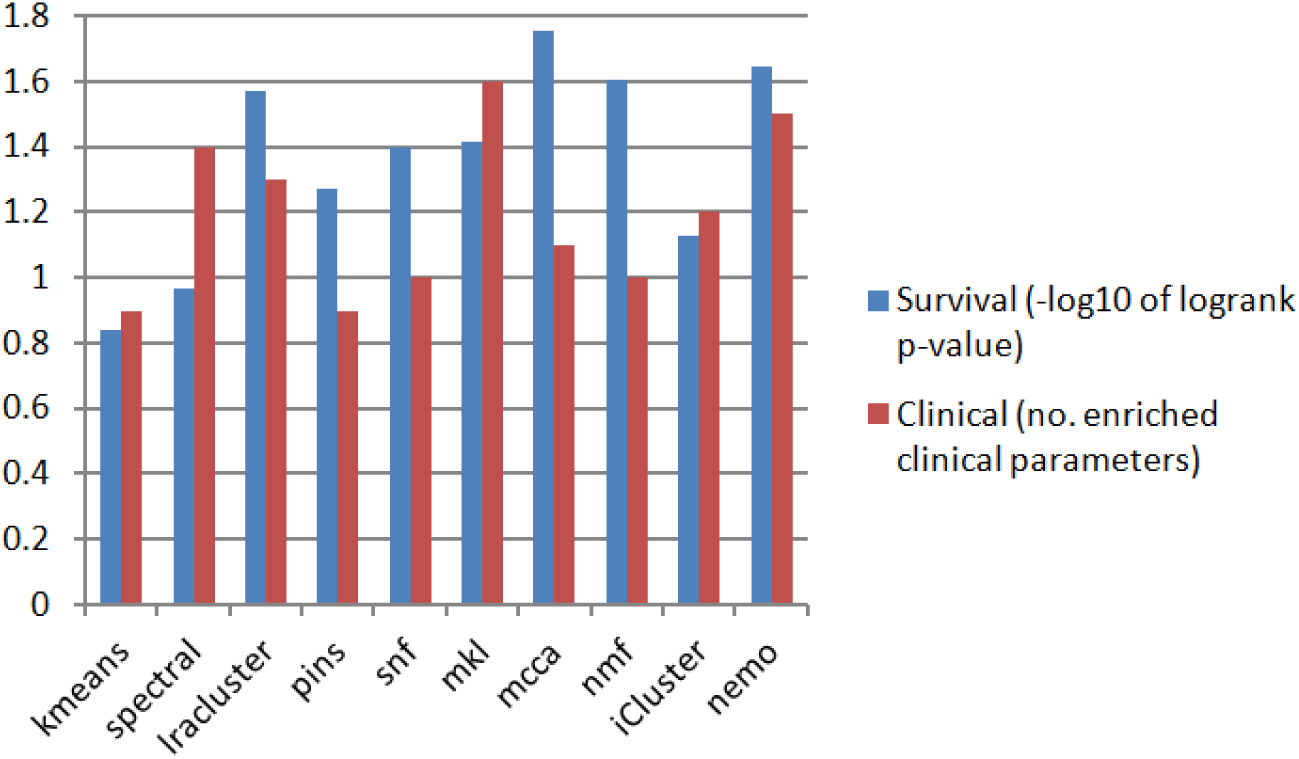
Mean performance of the ten algorithms on ten cancer datasets. Blue bars are the measured average differential survival between clusters (-log10 logrank test’s p-values). Red bars are the average number of enriched clinical parameters in the clusters.

Compared to the other methods, NEMO tended to choose an intermediate number of clusters per dataset (average 4.5). This number of clusters is small enough so that the clusters will be highly interpretable, but still capture the heterogeneity among cancer subtypes.

NEMO had the the second fastest runtime after spectral clustering of the concatenated omics matrix. All methods except for LRAcluster and iClusterBayes took only a few minutes to run on datasets with hundreds of samples and tens of thousands of features. However, due to NEMO’s simple integration step, it was the fastest of all non-trivial integration methods, including other similarity-based methods (SNF and rMKL-LPP). For rMKL-LPP, the time reported does not include the similarity computation, as this code was not provided by the authors, but was implemented by us, so its total runtime is higher. Details regarding the hardware used appear in Supplementary File 1. We note that since NEMO’s integrated network is sparse, its spectral clustering step can be further improved using methods for spectral clustering of sparse graphs (e.g, Lanczos, 1950). This advantage in runtime, and NEMO’s improved asymptotic runtime compared to other similarity-based methods (see Methods section) will become more important as the number of patients in medical datasets increases.

### Simulated partial datasets

We next evaluated NEMO’s performance on simulated partial datasets. We tested two scenarios. In the first we created two clusters using multivariate normal noise around the clusters’ centers, and then created two omics by adding to these data different normal noise for each omic (see Supplementary file 1). The simulation is therefore designed such that both omics share the same underlying clustering structure.

In the second scenario, we added a third omic that does not distinguish between the clusters. To simulate partial data, we removed the second omic data in an increasing fraction *θ* of randomly chosen samples, for *θ* ranging betwee 0 and 0.8. We generated 10 different full datasets, and for each dataset and for each value of *θ* we performed ten repeats. Here we report the average ARI between the computed and the correct clustering for each *θ*.

We compared NEMO’s performance to PVC, and also to MCCA and rMKL-LPP, the top performers on the full real data. To run PVC on the dataset, we subtracted the minimal observed value from each omic, making all values non-negative, and set PVC’s *λ* parameter to 0.01. Since PVC’s implementation supports only two omics, we ran it only in the first scenario. To run rMKL-LPP and MCCA on partial data, we completed the missing values using KNN imputation on the concatenated omics matrix. We set NEMO’s parameter *k* to half the number of samples as described in the Methods section. Full details about PVC’s execution appear in Supplementary File 1. MCCA was applied twice, using two low dimensional representations (See Supplementary File 1 for details).

Fig. 2 shows that NEMO outperformed other methods in both simulations. Furthermore, NEMO performed better on data that were not imputed than on data that were imputed. This shows the advantage of using NEMO directly on partial datasets, rather than performing imputation. In both scenarios, the performance of all methods deteriorated as the fraction of missing data increased. A notable exception was MCCA when using the low dimensional representation of the first omic. We believe this representation was barely affected by the second omic. Interestingly, adding a third omic that contributes no information to the clustering solution decreased the performance, but this decrease was minor for all methods.

**Figure 2:**
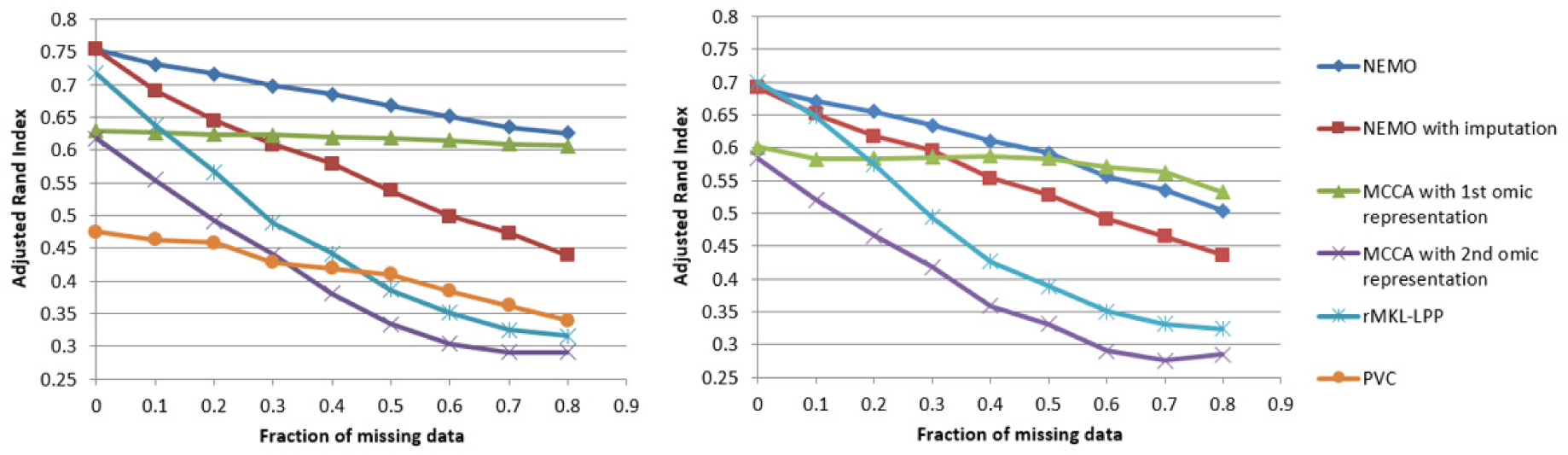
Performance on simulated partial data. We executed the algorithms with an increasing fraction of samples missing data in one of the omics, and compared the resulting clustering to the ground truth using ARI. The left plot uses two omics, and the right plot uses three omics where the third one contains only noise.

PVC performed poorly compared to NEMO in the two-omics simulation. In fact, PVC with all data for both omics performed worse than NEMO with 80% missing data in the second omic. We suspect that since PVC is based on linear dimension reduction, it does not capture the spheric structure of the clusters. Since PVC’s implementation supports only two omics, we could not test it in the second scenario.

### Partial cancer datasets

We next compared NEMO to other methods on partial cancer datasets, by simulating data loss on the ten full TCGA datasets analyzed above. We tested two scenarios, (1) using three omics for all subtypes, and (2) using only two omics, to allow comparison with PVC. We randomly sampled a fraction *θ* of the patients and removed their second omic data. The other omic(s) data were kept full. The *θ* values tested were between 0 and 0.7. In all datasets, the first omic was DNA methylation, and the second (from which samples were removed) was gene expression. In the three-omic scenario, the last omic was miRNA expression. We repeated each test five times, and the p-values reported here are the geometric averages of the observed p-values.

Full details on how each method was executed are in Supplementary File 1. We set the number of clusters in PVC to be the same as determined by NEMO, since no method to determine the number of clusters was suggested for PVC. We used survival analysis and enrichment of clinical labels to assess the quality of the clustering solutions. Full results for this analysis are in Supplementary File 3.

Figure 3 shows the mean results on three omics across all ten cancer types. NEMO performed best with respect to survival, followed by NEMO with imputation. rMKL-LPP performed best with respect to clinical parameters, followed by NEMO with and without imputation.

**Figure 3:**
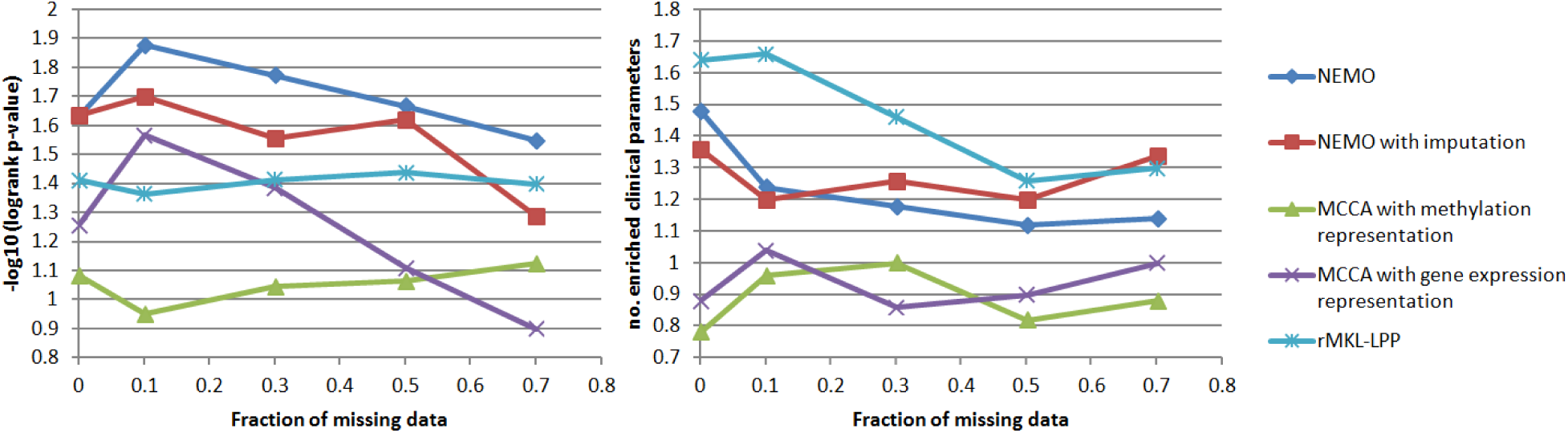
Performance on partial cancer datasets as a function of the fraction of samples missing data in one of the omics. Left: survival analysis. Right: clinical parameters. Results are averages across ten three-omics cancer datasets.

Note that in contrast to simulated data, here the performance of the methods did not consistently deteriorate as more data were removed. This is somewhat surprising, as gene expression (the omic that was partially removed) is believed to be the most informative omic. The difference between the performance of MCCA for full data here (*θ* = 0) and its previous results (Figure 1) is due to the fact that MCCA optimizes its objective with respect to one omic at a time, which makes the solution sensitive to the order of the omics.

In the second scenario, out of the datasets that had statistically significant survival results, NEMO was best performer for AML, GBM and SARC, while PVC was best for BIC and SKCM (Supplementary Figure 3). PVC (using the number of clusters determined by NEMO) had best mean survival and clinical enrichment across all datasets (Supplementary Figure 5). This shows the merit of PVC (and of NEMO’s method to determine the number of clusters) in datasets with two omics. Interestingly, for both NEMO and PVC, the mean performance across all ten full two-omics datasets was better than the performance of all methods on full three-omics datasets in terms of survival (Figure 3 and Supplementary Figure 5). Performing imputation increases the runtime of the algorithms. For example, the average time (across 5 runs) to perform imputation for the BIC dataset with methylation and mRNA expression omics, when *θ* = 0.5, was 560 seconds. This is a necessary preprocessing step for methods that do not directly support missing data. In contrast, not only do NEMO and PVC not require imputation, but they also run faster as the fraction of missing data increases. The runtime of NEMO on the same two-omic BIC dataset decreased from 42 seconds with full data to 21 seconds with *θ* = 0.7. PVC was slower than NEMO, and its runtime decreased from 92 seconds to 27 seconds.

### Robustness analysis

We sought to assess NEMO’s robustness with respect to the parameter *k*, the number of neighbors. We performed clustering on the ten cancer datasets using *k* = 25, 35, …, 105, a range that includes all *k* values we used in the full and partial data analyses. Supplementary Fig. 6 shows the p-values for logrank test for each value of *k*. Generally, the performance of NEMO was robust with respect to *k*. In a few cases, such as the GBM, SKCM and SARC datasets, the results varied more depending on *k*. This is partially explained by the different number of clusters NEMO chose for different values of *k* (see Supplementary Figure 7). We conclude that usually changing *k* has little effect on the number of clusters and on the significance. In those cases where significance changed with *k*, it was usually a result of change in the number of clusters chosen (compare Supplementary Figures 6, 7, and 8).

### Acute Myeloid Leukemia analysis

We applied NEMO to an AML cohort of 197 patients from TCGA. This is a partial dataset, containing 173 patients with gene expression profiles, 194 with methylation and 188 with miRNA profiles. As it is partial, the dataset cannot be directly clustered using other algorithms for multi-omics clustering. To apply these methods, one must limit analysis to a sub-cohort of 170 patients that have full data, or perform imputation. NEMO suggested five clusters for this dataset. When plotting survival curves of the clusters (Fig. 4), we found them to be significantly different (p-value = 3.5e-4). The significance was higher than obtained by all other nine clustering methods on the full data subcohort (lowest p-value 1.3e-3 using k-means). This shows the higher significance gained from analyzing more samples, including partial data.

**Figure 4:**
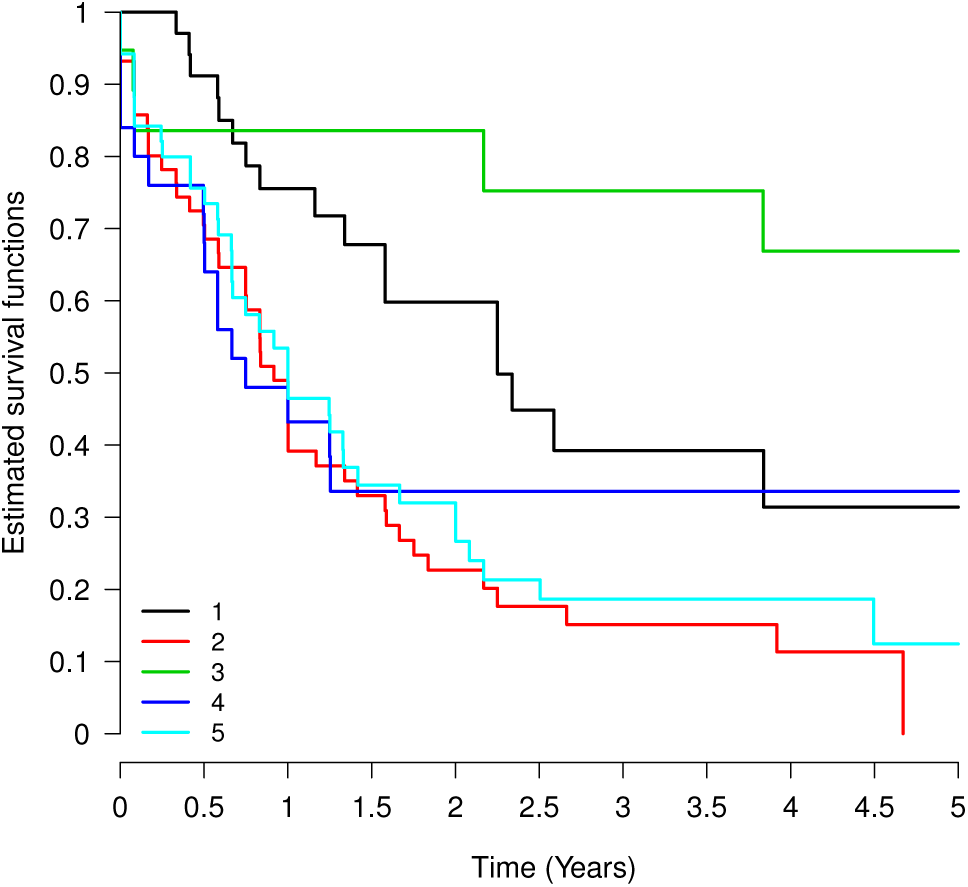
Kaplan-Meier plot for the five clusters obtained by NEMO on the AML partial dataset (logrank p-value=3.5e-4).

We compared the prognostic value of NEMO’s clusters to that of the FAB (French-American-British) classification. FAB is a well-accepted clinical classification for AML tumors (Bennett *et al*., 1976), which is based on quantification of blood cells. We performed logrank test using the FAB label as clustering solution, and obtained a p-value 5.4e-2, which shows NEMO’s favorable prognostic value. Executing NEMO using only a single omic, results for gene expression, methylation and miRNA expression data had logrank p-values 3.4e-2, 3.4e-3 and 3.7e-3 respectively. These results demonstrate the improved clustering obtained by NEMO using multi-omic data.

We performed enrichment analysis for each NEMO cluster using the PROMO tool, which allows systematic interrogation of all clinical labels (Netanely *et al*., 2016). In addition to the significantly differential survival, the clusters were found to be enriched in other clinical labels. Cluster 1 had particularly young patients, and showed favorable prognosis. Cluster 2 had poor prognosis, older patients, and was enriched with FAB label ”M0 undifferentiated”. 17 out of 19 patients with label ”M0 undifferentiated” appeared in this cluster. This label corresponds to the undifferentiated acute myeloblastic AML subtype, which is known to have poor prognosis (Bene *et al*., 2001). Cluster 3 showed favorable prognosis, and was enriched with the M3 FAB label, which corresponds to the acute promyelocytic leukemia (APL) subtype. All 19 patients in this cluster were labeled with M3, and only one patient outside cluster 3 had this label. APL is caused by a translocation between the genes RARA on chromosome 17 and PML on chromosome 15, and is known to have favorable prognosis (Wang and Chen, 2008). Cluster 4 was enriched with the M5 label, which corresponds to acute monocytic leukemia. Indeed, it was also enriched with a high monocyte count. Finally, cluster 5 was enriched with patients with no known genetic aberrations. All the clustering results and enriched clinical labels are included in Supplementary Files 4 and 5.

## Discussion

We presented the NEMO algorithm for multi-omics clustering, and tested it extensively on cancer datasets and in simulation. NEMO is much simpler than existing multi-omics clustering algorithms, has improved performance on full datasets, improved performance on partial datasets without requiring missing data imputation, and faster execution.

NEMO’s simplicity makes it more flexible and more easily adapted to different circumstances. It requires only the definition of a distance between two samples within an omic, and can therefore support additional omics, numerical, discrete and ordinal features, as well as more complicated feature types, such as imaging, EMR data and microbiome. In addition to enabling clustering, the network produced by NEMO represents the similarity between samples across all omics, and can thus be used for additional computational tasks. Future work will test the usability of NEMO on discrete data types, and of its output network for tasks other than clustering.

We showed that NEMO can be used to analyze partial multi-omic datasets, i.e. ones in which some samples lack measurements for all omics. Partial datasets are ubiquitous in biology and medicine, and methods that analyze such datasets hold great potential. This challenge is exacerbated by the high cost of high-throughput experiments. While the price of some experiments is decreasing, it is still high for other omics. Methods that analyze partial datasets may affect experimental design and reduce costs, and, as we demonstrated, they can outperform full-data methods applied only to the subset of samples that have all omics. The demand for algorithms that analyze partial datasets is likely to further increase, as more high throughput methods become prevalent, and the number of omics in biomedical datasets increases.

NEMO has several limitations. First, in partial data, each pair of samples must have at least one omic in common. This assumption holds if one omic was measured for all patients, which is often the case for gene expression. Second, while we showed that NEMO is robust to the choice of *k*, the number of neighbors, that choice requires further study. Third, unlike some dimension reduction methods, NEMO does not readily provide insight on feature importance. Given a clustering solution, importance of features to clusters can be computed using differential analysis.

We compared NEMO to PVC in the context of missing data. PVC was developed within the machine learning community for the task of partial multi-view clustering, and has not been applied to omic data previously. Remarkably, on average, in terms of survival analysis, PVC (using the number of clusters of NEMO) outperformed all other methods on the partial cancer datasets with two omics, while NEMO was better on the simulated partial datasets. As PVC is limited to two omics, extension of that NMF-based algorithm to more omics and to include a mechanism for determining the number of clusters is desirable.

In some of the cancer datasets the results obtained using only mRNA expression and DNA methylation were superior to those achieved when also considering miRNA expression. In addition, we did not observe a significant decrease in performance when removing a fraction of the gene expression data for cancer patients. This phenomenon suggests that multi-omics clustering does not necessarily improve with more omics (see also Rappoport and Shamir 2018). While NEMO performed well with an additional omic that contains no signal, future work is needed to deal with omics that contain noise or contradicting signals.

## Conclusion

Clustering cancer patients into subgroups has the potential to define new disease subtypes that can be used for personalized diagnosis and therapy. The increasing diversity of omics data as well as their reduced cost creates an opportunity to use multi-omic data to discover such subgroups. NEMO’s simplicity, efficiency and efficacy on both full and partial datasets make it a valuable method for this challenge.

## Acknowledgements

The results published here are based upon data generated by The Cancer Genome Atlas managed by the NCI and NHGRI. Information about TCGA can be found at http://cancergenome.nih.gov. The contribution of N.R. is part of Ph.D. thesis research conducted at Tel Aviv University.

